# Deep Mutational Scanning Identifies Sites in Influenza Nucleoprotein That Affect Viral Inhibition by MxA

**DOI:** 10.1101/071969

**Authors:** Orr Ashenberg, Jai Padmakumar, Michael B. Doud, Jesse D. Bloom

**Affiliations:** Division of Basic Sciences and Computational Biology Program, Fred Hutchinson Cancer Research Center, Seattle, WA, USA; Department of Genome Sciences, University of Washington, Seattle, WA, USA; Medical Scientist Training Program, University of Washington School of Medicine, Seattle, WA, USA

## Abstract

The innate-immune restriction factor MxA inhibits influenza replication by targeting the viral nucleoprotein (NP). Human influenza is more resistant than avian influenza to inhibition by human MxA, and prior work has compared human and avian viral strains to identify amino-acid differences in NP that affect sensitivity to MxA. However, this strategy is limited to identifying sites in NP where mutations that affect MxA sensitivity have fixed during the small number of documented zoonotic transmissions of influenza to humans. Here we use an unbiased deep mutational scanning approach to quantify how all ≈10,000 amino-acid mutations to NP affect MxA sensitivity. We both identify new sites in NP where mutations affect MxA resistance and re-identify mutations known to have increased MxA resistance during historical adaptations of influenza to humans. Most of the sites where mutations have the greatest effect are almost completely conserved across all influenza A viruses, and the amino acids at these sites confer relatively high resistance to MxA. These sites cluster in regions of NP that appear to be important for its recognition by MxA. Overall, our work systematically identifies the sites in influenza nucleoprotein where mutations affect sensitivity to MxA. We also demonstrate a powerful new strategy for identifying regions of viral proteins that affect interactions with host factors.

**Author Summary:** During viral infection, human cells express proteins that can restrict virus replication. However, in many cases it remains unclear what determines the sensitivity of a given viral strain to a particular restriction factor. Here we use a high-throughput approach to measure how all amino-acid mutations to the nucleoprotein of influenza virus affect restriction by the human protein MxA. We find several dozen sites where mutations substantially affect influenza’s sensitivity to MxA. While a few of these sites are known to have fixed mutations during past adaptations of influenza to humans, most of the sites are broadly conserved across all influenza strains and have never previously been described as affecting MxA resistance. Our results therefore show that the known historical evolution of influenza has only involved substitutions at a small fraction of the sites where mutations can in principle affect MxA resistance. We suggest that this is because many sites are already broadly fixed at amino acids that confer high resistance.

## Introduction

Influenza proteins must evade immunity while maintaining their ability to function and interact with host cell factors [1]. The effects of immune pressure on influenza evolution have been most studied in the context of adaptive immunity, with numerous studies showing how antibodies and T-cells drive the fixation of immune-escape mutations in viral proteins [2–5]. However, the innate immune system also exerts selection on influenza via the interferon-stimulated expression of restriction factors, some of which target viral proteins and inhibit their function. The first anti-influenza restriction factor to be discovered, the murine protein Mx1, was initially described over 50 years ago [6–8]. It is now known that Mx1 and its human ortholog MxA inhibit influenza by interacting with the viral nucleoprotein (NP) [9–14]. However, the exact mechanistic details of the inhibitory interaction between MxA and NP remain incompletely understood.

Influenza counteracts the inhibitory effects of MxA through two distinct strategies: it generally blocks the interferon-response that drives expression of MxA [15], and it fixes specific amino-acid mutations in NP that reduce its sensitivity to MxA [16, 17]. The importance of the second of these two strategies has been elegantly demonstrated by studies comparing the MxA sensitivity of different viral strains. NPs from avian and swine influenza are more sensitive to human MxA than NPs from human influenza [16, 17]. NPs from non-human influenza have been introduced into circulating human influenza strains twice over the last century: once in 2009 from swine influenza [18], and once in or before 1918 probably from avian influenza [19–21]. By functionally characterizing the effects of mutations at sites that differ between these human influenza NPs and their predecessors from non-human viral strains, Mänz et al [17] identified a small set of sites in NP where mutations affect MxA resistance. Riegger et al [22] subsequently identified another site in NP where a mutation has increased the MxA resistance of an avian H7N9 virus that has undergone numerous non-sustained zoonotic transmissions to humans.

Characterizing sites that differ between non-human and human influenza strains is a powerful strategy to identify mutations that have historically contributed to the adaptation of NP to avoid recognition by MxA. However, it is an incomplete approach for mapping the full set of sites in NP that affect sensitivity to MxA. Evolution is stochastic [23–25], meaning that any given adaptation event will sample only some of the possible mutations that confer MxA resistance. In addition, adaptation of non-human influenza to humans only favors mutations at NP sites that initially encode an amino acid that confers relatively high sensitivity to MxA. Sites at which avian and swine influenza already possess MxA-resistant amino acids will not be identified by cross-species comparison. Therefore, more systematic approaches are needed to fully characterize the sites in NP that affect MxA resistance.

Systematic measurement of how all amino-acid mutations affect a protein phenotype has recently become possible with the advent of deep mutational scanning [26, 27]. This massively parallel experimental technique involves generating a library of mutants, imposing a functional selection, and using deep sequencing to determine the frequency of each mutation before and after selection. Deep mutational scanning has already been used to examine the functional effects of most mutations to several influenza proteins [28–35].

Here we use deep mutational scanning to quantify how every amino-acid mutation to the NP of a human influenza strain affects sensitivity to MxA. This unbiased approach enables us to identify mutations that both increase and decrease MxA sensitivity. Therefore, in addition to re-identifying some sites where mutations have previously been shown to adapt influenza to MxA, we are able to identify new sites that affect MxA sensitivity. We find that mutations at NP site 51 have the largest effect on MxA sensitivity, and that almost all known influenza A strains already possess an amino acid at this site that confers high resistance. Overall, our work finds new sites affecting MxA resistance that could not have been identified by comparing viral strains across species, and introduces a framework for comprehensively profiling the effect of all mutations to viral proteins on recognition by restriction factors.

## Results

### A deep mutational scan for NP mutations that affect MxA resistance

Our goal is to understand which sites in influenza NP determine its sensitivity to MxA. We can do this by experimentally quantifying how all amino-acid mutations to NP affect sensitivity to MxA. Systematic measurements of this type can be made using the deep mutational scanning approach outlined in Fig 1. This approach involves creating influenza viruses that carry a diverse set of NP mutations, growing these viruses in cells that do or do not express human MxA, and then using deep sequencing to identify mutations that are enriched or depleted in one condition versus the other. Mutations that are enriched in MxA-expressing cells relative to control cells increase MxA resistance, whereas mutations that are relatively depleted in MxA-expressing cells increase MxA sensitivity.

**Fig 1.**
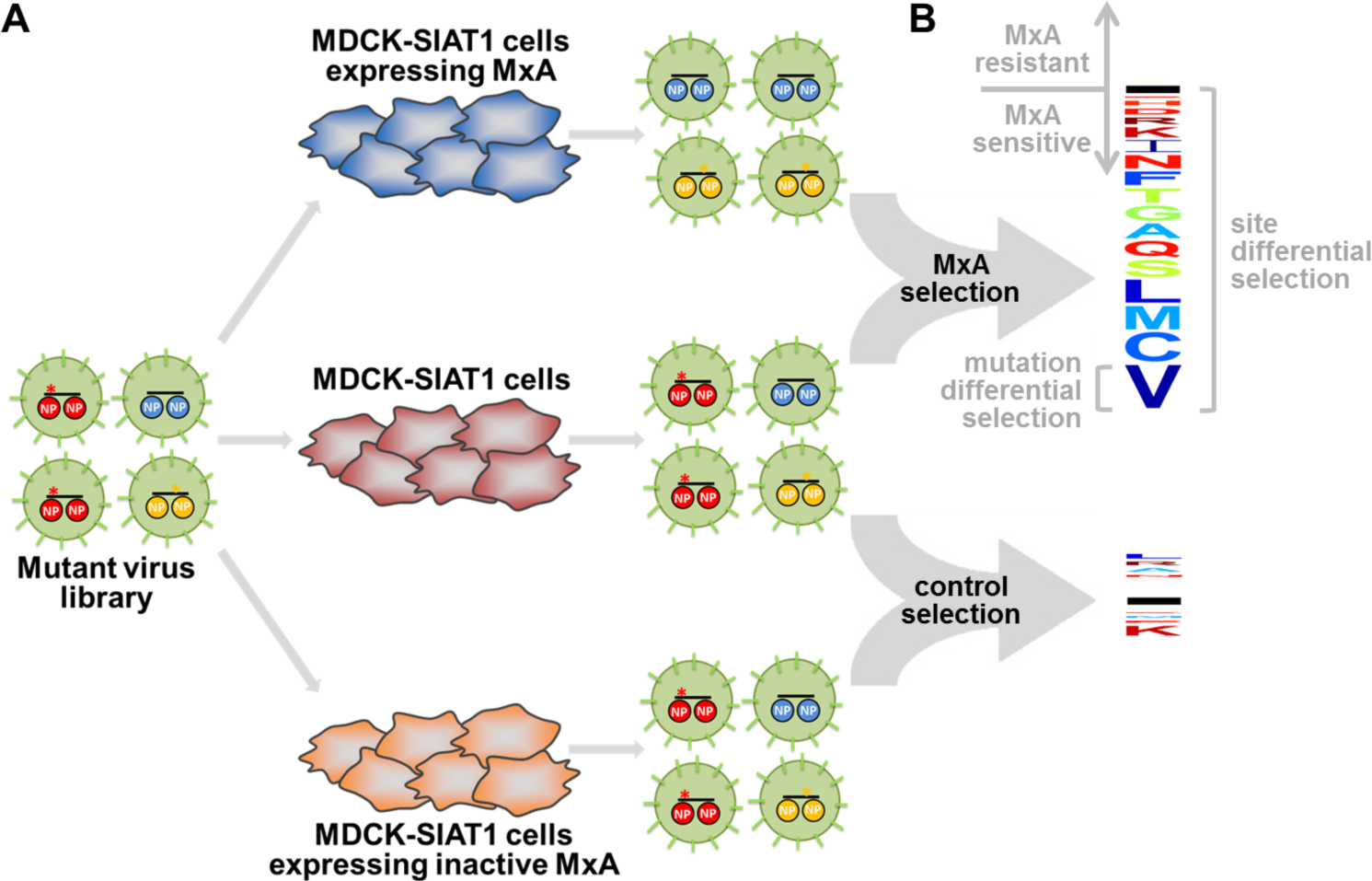
A deep mutational scan for NP mutations that affect MxA resistance. (**A**) We created a library of influenza variants carrying all viable amino-acid mutants of NP. We infected cells expressing human MxA, not expressing human MxA, or expressing inactive MxA, and used deep sequencing to quantify the enrichment or depletion of each mutation in each condition. (**B**) The effect of each mutation on MxA sensitivity is computed as the logarithm of its relative frequency in MxA-expressing versus non-expressing cells. These measurements are summarized in the logo plots, where letters above the black horizontal line represent amino acids that increase MxA resistance, and letters below the line represent amino acids that increase MxA sensitivity. The overall differential selection at a site is the total height of the letter stack. Letters are colored according to hydrophobicity. The cells expressing the inactive MxA provide a control to estimate experimental noise. At the example site shown (site 283), most mutations increase MxA sensitivity, and the actual differential selection in the MxA-expressing cells is much greater than the noise measured in the cells expressing inactive MxA.

We chose to perform our deep mutational scan on a NP from a human-adapted influenza strain, A/Aichi/2/1968 (H3N2). We reasoned that use of a human-adapted NP should allow us to better detect mutations that decrease MxA resistance, as well as identify any resistance-enhancing mutations that have not already fixed in human influenza. The use of a human-adapted NP makes our approach complementary to previous studies that have focused on mutations that increase the MxA resistance of non-human strains of influenza [17, 22]. We have previously described duplicate libraries of influenza viruses that carry nearly all amino-acid mutations to the Aichi/1968 NP that are compatible with viral growth [32]; these libraries formed the starting point for the work performed here.

During viral infection of normal human cells, MxA expression is induced by activation of the interferon response, which varies from cell to cell [36, 37]. But our controlled experiment (Fig. 1) requires cells that never or always express a functional human MxA. For our MxA-deficient cells, we chose MDCK-SIAT1 cells, a variant of the Madin Darby canine kidney cell line. We chose these cells for two reasons. First, the canine MxA ortholog lacks anti-influenza activity against all influenza strains tested to date [38, 39]. Second, MDCK-SIAT1 cells support robust growth of influenza, and are therefore well-suited to maintaining the diversity of our virus library. To create MxA-expressing cells, we engineered a MDCK-SIAT1 cell line to constitutively expresses human MxA. We also created a control cell line that constitutively expresses the T103A mutant of MxA, which is inactive against influenza [40]. For both cell lines, we verified MxA protein expression (Fig 2A). We also verified that constitutive expression of wildtype but not T103A MxA profoundly inhibits viral replication (Fig 2B). This inhibition demonstrates that even human-adapted influenza NP is sensitive to sufficiently high levels of MxA.

**Fig 2.**
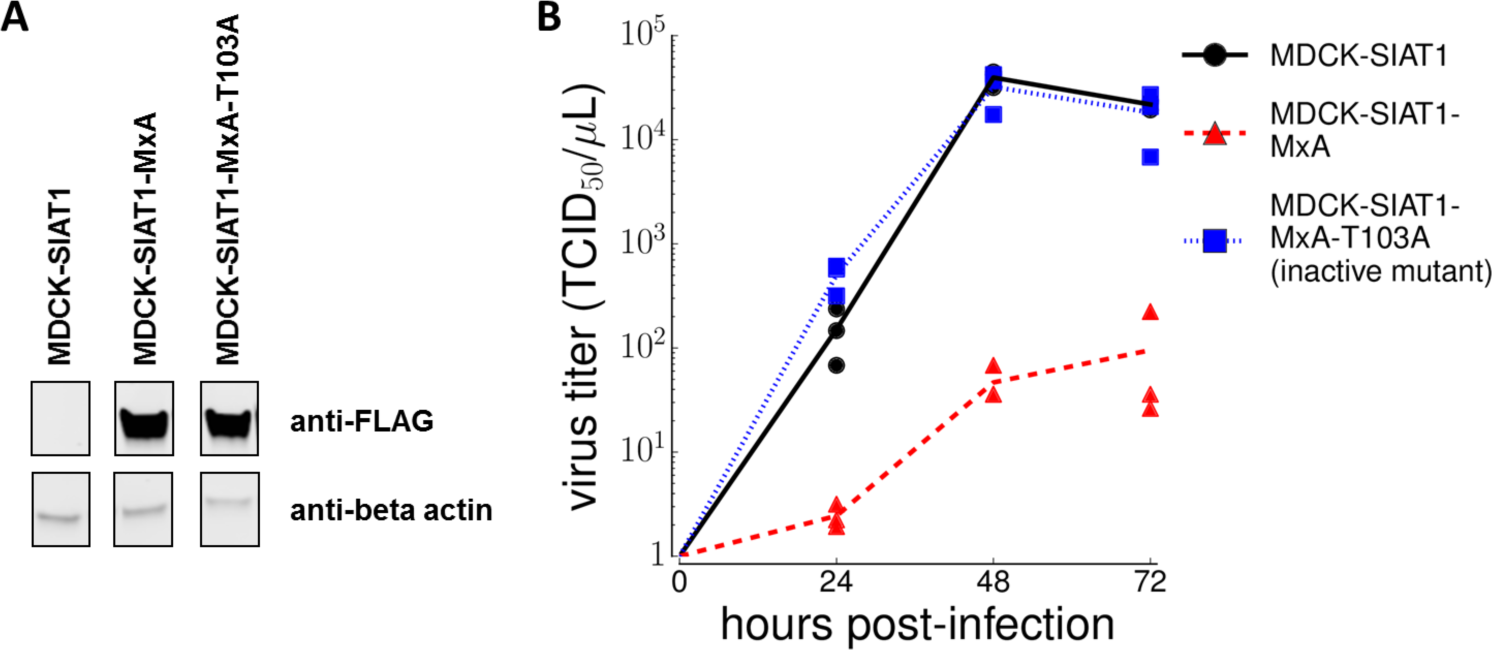
Viral growth is inhibited by MxA. (**A**) We created MDCK-SIAT1 cells expressing FLAG-tagged MxA, and verified MxA expression by Western blot. (**B**) We infected the cell lines with virus carrying the A/Aichi/2/1968 NP at an MOI of 0.01, and measured viral titers every 24 hrs. Viral growth was inhibited in MxA-expressing cells, but not in non-expressing cells or cells expressing MxA with the T103A inactivating mutation.

We then infected our virus libraries into all three cell lines as indicated in Fig 1A. In order to maintain the diversity of the libraries, we infected each cell line with 5 × 10^6^ TCID_50_ of virus. We performed the infections at a multiplicity of infection (MOI) of 0.1 to reduce viral co-infection and subsequent genetic complementation. We then isolated viral RNA from infected cells after 48 hours, and deep sequenced NP to determine the frequency of each mutation in each selective condition. We used overlapping paired-end Illumina reads to reduce the sequencing error rate (S1 Fig shows that this strategy reduced the net rates of errors associated with sequencing, PCR, and reverse transcription to below the frequency of the actual mutations of interest). We performed this experiment independently for each NP virus library, meaning that all high-throughput measurements were made in true biological duplicate. Our expectation was that analyzing the deep sequencing data would enable us to identify mutations that affect MxA resistance as shown in Fig 1B.

### Analysis of the deep mutational scanning data identifies sites where mutations affect MxA sensitivity

We estimated the effect of each mutation on MxA resistance by computing the logarithm of its frequency in the MxA-expressing cells relative to the non-expressing cells. We refer to this quantity as the mutation differential selection. We estimated the total differential selection at each site by summing the absolute values of the differential selection on each mutation at the site. Fig 1B graphically illustrates our measures of mutation and site differential selection. Fig 3A shows that our two independent replicates of deep mutational scanning yielded reproducible estimates of the differential selection at each site. The estimates of the differential selection on individual mutations were also significantly correlated between replicates, although they were noisier than the per-site ones (S2 Fig).

**Fig 3.**
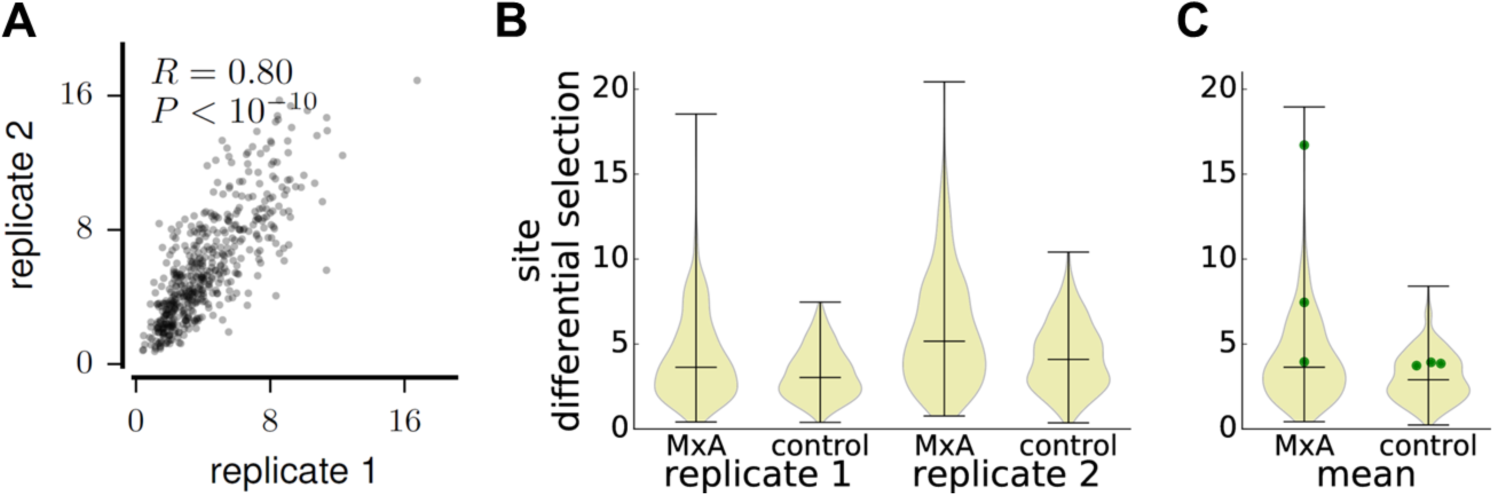
MxA exerts strong selection at some sites in NP. (**A**) The correlation between site differential selections measured by the two independent experimental replicates. Estimates of the differential selection from MxA at each site are highly reproducible. (**B**) The distributions of differential selections across all sites in NP in the actual MxA selections and the control selections with inactive MxA. The distributions are shown as violin plots, with the three horizontal bars marking the minimum, median, and maximum of the site differential selections. For both replicates, the MxA selections have a small number of sites where MxA selection greatly exceeds the background measured in the control selection. (**C**) The distributions of site differential selections averaged across the replicates. The three sites (100, 283, and 313) shown by [17] to affect the MxA sensitivity of the 1918 virus are shown as green points. All three sites have above-median effects on MxA sensitivity in our MxA selection experiments.

One way to test if the sensitivity of our experiments exceeded their noise is to compare the magnitudes of the differential selections observed in the actual selection with MxA-expressing cells versus the control selection with cells expressing inactive MxA (Fig. 1A). Fig. 3B shows the distribution of differential selection values across all sites as estimated in the MxA and control selections for each replicate. For each replicate, there was a long tail of sites with strong differential selection in the MxA selection that exceeded any value observed in the control selection with inactive MxA. S2 Fig shows that similar results are obtained when examining differential selection at the level of mutations rather than sites. Therefore, at a subset of sites, MxA exerts selection that substantially exceeds the background noise of the experiments.

We next tested whether our results were consistent with existing knowledge about how mutations to NP affect MxA resistance. For this test and the remainder of this paper, we use the average of the measurements from the two replicates (S3 Fig shows these average measurements for all sites.) We examined the three NP mutations previously shown to be mostly responsible for the increased MxA resistance of the human 1918 pandemic H1N1 strain relative to its its avian influenza ancestors (according to [17], these are R100V, L283P, and F313Y). The Aichi/1968 NP in our study is a descendant of this 1918 NP and retains all three MxA-resistance mutations. We therefore expected that reverting these mutations would increase MxA sensitivity, barring effects due to changes in NP sequence context between 1918 and 1968. Consistent with this expectation, the differential selection for reverting each mutation to its consensus identity in avian NP was negative (S2 Fig). All three mutations also occurred at sites that had differential selection that exceeded the median (Fig 3C). These results show that despite a half-century of sequence divergence, the sites of the mutations that conferred MxA resistance on the 1918 virus remain important determinants of this phenotype in the NP used in our study.

We next sought to identify the NP sites that most affected MxA resistance. There were 29 sites where the differential selection from MxA exceeded the background noise in the control selection. Fig 4A shows these 29 sites ranked by their differential selection values. Site 283, which is one of the sites most responsible for the MxA resistance of the 1918 pandemic virus [17], ranks second in our data (Fig 4A). But most sites predicted by our deep mutational scan to have the largest effects on MxA resistance have not previously been described as impacting this phenotype. Interestingly, at all these sites, the greatest differential selection is from mutations away from the wildtype amino acid that increase MxA sensitivity.

**Fig 4.**
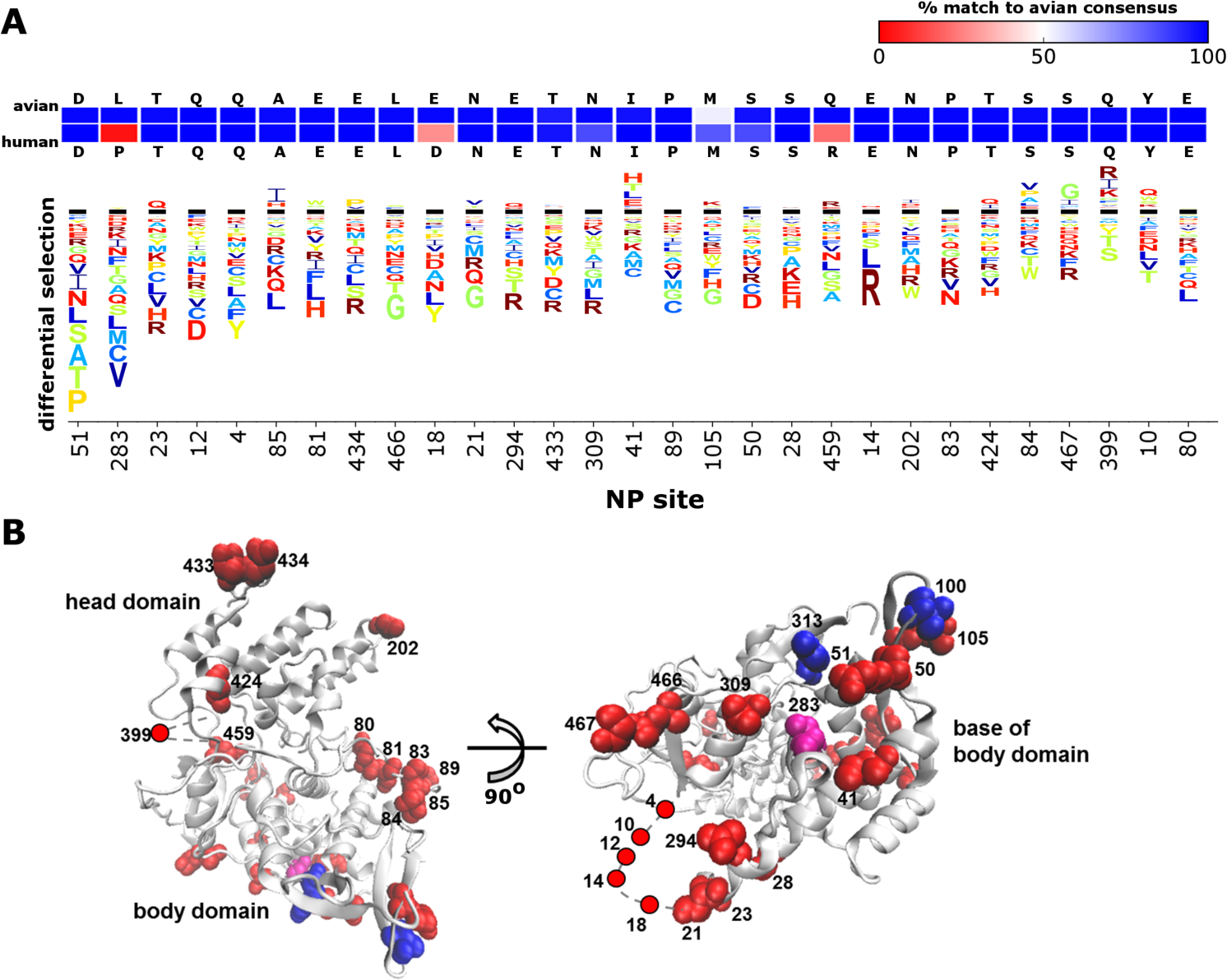
The sites with the greatest effect on MxA sensitivity in our experiments. (**A**) Differential selection from MxA for the 29 sites where total differential selection exceeds the maximum in the control selection. The logo stacks have the same meaning as in Fig 1B. The sites are ordered by total differential selection. Above the logo stacks are the avian and human consensus residues for each site. The two rows of boxes are color coded according to the percent of avian (top row) or human (bottom row) NP sequences that match the avian consensus at that site. The colors range from blue (100% match to avian consensus) to red (0% match to avian consensus). (**B**) The 29 sites mapped onto NP’s structure (PDB 3ZDP) as red spheres. Most sites strongly affecting MxA sensitivity are at the base of the body domain, in solvent-exposed loops, or near the N-terminus. The N-terminus and a loop (residues 392 to 407) are unresolved in the structure and so are modeled with a gray dashed line. The sites previously identified [17] as being responsible for the MxA resistance of the 1918 virus are shown as blue spheres. Site 283 is in both the 1918-resistance set and our set of 29 sites, and so is shown in purple.

Strikingly, 26 of the 29 sites where mutations most influence MxA resistance have the same consensus amino acid in avian and human influenza NP sequences (Fig 4A). Therefore, although these sites appear to be important determinants of the interaction between NP and MxA, they have not undergone extensive substitution during the adaptation of influenza to humans–presumably because they already possess an amino acid that confers resistance. The broad conservation of these sites also explains why they have not previously been identified by studies examining mutations that fixed during recent influenza evolution in nature.

The sites that most affected MxA resistance are on the surface of the monomeric structure of NP (Fig 4B). Twelve of these sites clustered at the base of the NP body domain, which is also the location of the three mutations that contributed to the MxA resistance of the 1918 virus [17]. The remaining sites mostly clustered either in a flexible basic loop known to affect RNA binding (residues 73 to 91) or in the N-terminus of NP, which is disordered [41–44]. Overall this structural mapping reinforces the central importance of the base of the NP body domain to MxA sensitivity, but shows that other NP surface regions may also contribute.

### NP site 51 is a major determinant of MxA sensitivity

The two NP sites with the greatest differential selection were 51 and 283 (Fig 4A). The mutation L283P has previously been characterized as increasing the MxA resistance of the 1918 virus relative to its avian ancestor [17], and our high-throughput data concur that mutating this site to the avian identity (or indeed to almost any amino acid other than P) substantially increases MxA sensitivity (Fig 4A, S4 Fig). But while site 283 has clearly undergone important change during influenza evolution, site 51 is almost completely conserved as D across all NPs from human, swine, equine, and avian influenza A strains (S1 Table). Our high-throughput data suggest that mutating site 51 to most other amino acids should greatly increase MxA sensitivity. Structurally, site 51 is located near site 283 (Fig 4B), and is adjacent to sites 52 and 53, where mutations have been shown to affect the MxA resistance of the 2009 pandemic H1N1 [17] and H7N9 [22] strains, respectively. Interestingly, a mutation at site 51 that increased MxA sensitivity arose as a secondary change in an avian influenza virus that was engineered for increased MxA resistance [45].

To validate the finding of our high-throughput experiments that site 51 is a major determinant of MxA sensitivity, we engineered variants of the Aichi/1968 NP carrying a variety of mutations at this site. We selected five amino-acid mutations that our high-throughput data suggest should reduce MxA resistance by varying degrees (S5 Fig). As a control, we also designed a synonymous mutation at site 51 (D51Dsyn) that was not expected to affect MxA sensitivity. To test these mutations, we measured the effect of each mutation on polymerase activity in the presence and absence of MxA. Active influenza polymerase can be reconstituted in cells, and this polymerase activity is sensitive to inhibition of NP by MxA [9]. We expected that polymerase activity would be more inhibited with mutant NPs that had increased MxA sensitivity. In the absence of MxA, the D51Dsyn mutation had similar polymerase activity to the wildtype NP while all five amino-acid mutations modestly decreased polymerase activity (Fig 5A). We compared these activities in the absence of MxA to those measured in cells expressing MxA, and quantified the effect of each mutation on MxA resistance by dividing its activity in the presence of MxA by its activity in the absence of MxA. The wildtype NP and the D51Dsyn mutant were slightly inhibited by MxA, with activity decreasing to ~80% of its original value (Fig 5B). As predicted by our high-throughput deep mutational scanning, all five amino-acid mutants at site 51 were more strongly inhibited, with activity decreasing to between 24% and 54% of its original value (Fig 5B). This result indicates that multiple different mutations away from D at site 51 substantially increase MxA sensitivity as measured by polymerase activity.

**Fig 5.**
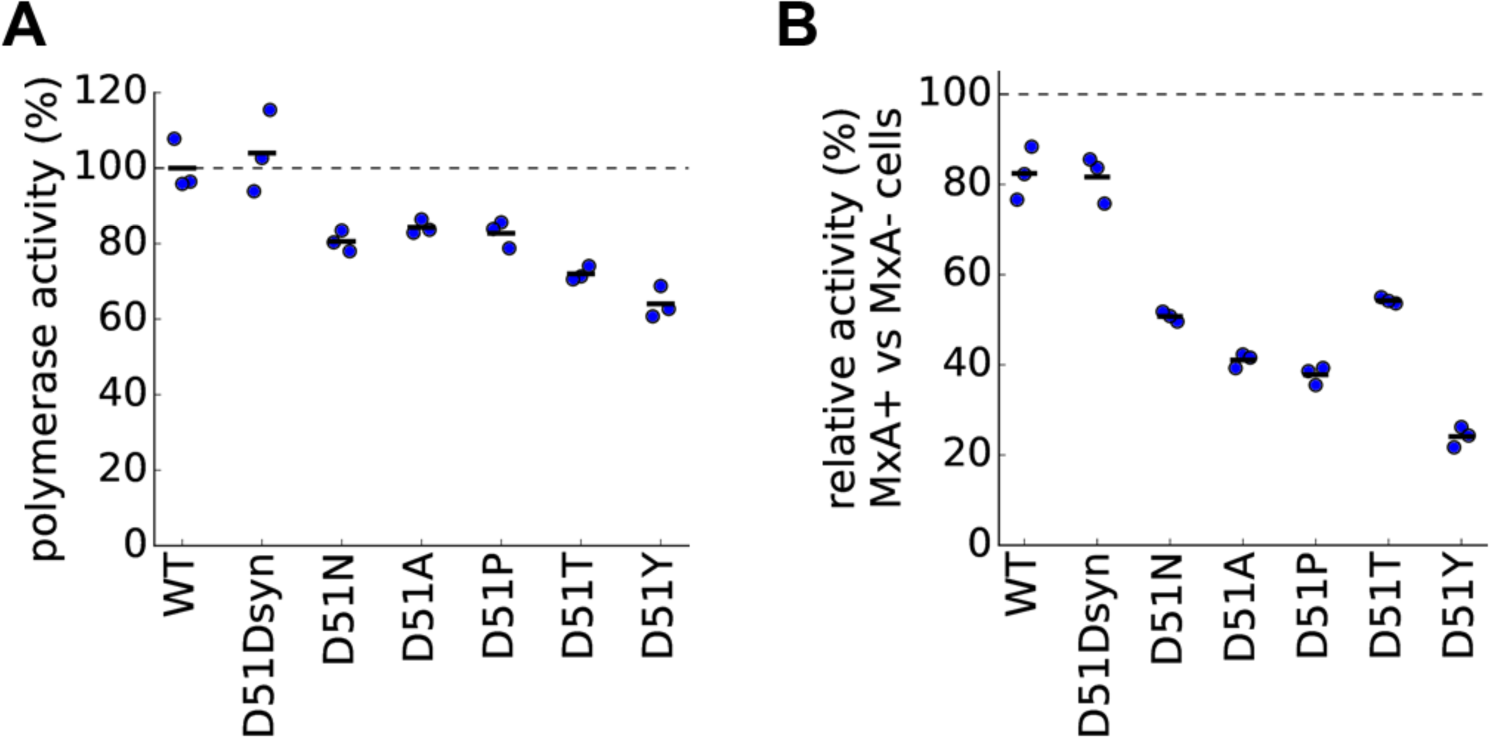
Mutations to site 51 in NP increase MxA sensitivity as measured by polymerase activity. (**A**) To measure polymerase activity, we transfected unmodified MDCK-SIAT1 cells not expressing MxA with plasmids for NP and the other polymerase-complex proteins as well as a GFP reporter viral RNA. The plot shows the levels of the GFP reporter for each mutant NP relative to wildtype NP, which is set to 100%. (**B**) To measure the change in MxA sensitivity for each mutation, we transfected MxA-expressing and non-expressing cells with the same plasmids as in (A). For each mutant, the plot shows the levels of the GFP reporter in MxA-expressing cells normalized by the same mutant’s activity in non-expressing cells.

To confirm that the decreased MxA resistance of polymerase activity correlated with the effect of MxA on viral replication, we carried out competition experiments between wildtype and mutant viruses. Such competition experiments provide a sensitive and internally controlled way to measure the relative fitness of two viral variants. We used reverse genetics to generate influenza viruses carrying wildtype NP, the D51Dsyn mutation, or the D51N mutation (Fig. 6A). We mixed each mutant virus with wildtype virus at a 1:1 ratio of infectious particles, and infected MxA-expressing or non-expressing cells at a low MOI. At 10 and 54 hours post-infection, we isolated viral RNA and determined the frequency of each variant by deep sequencing. As expected, the wildtype D51 variant greatly increased in frequency relative to the MxA-sensitive D51N mutant in MxA-expressing cells, whereas the two variants remained at similar frequencies in cells not expressing MxA (Fig 6B). Also as expected, the wildtype D51 variant and its synonymous variant remained at roughly equal frequencies in the control competitions (Fig 6B). This competition experiment verifies that an amino-acid mutation away from the wildtype identity at site 51 strongly increases MxA sensitivity as measured by viral growth.

**Fig 6.**
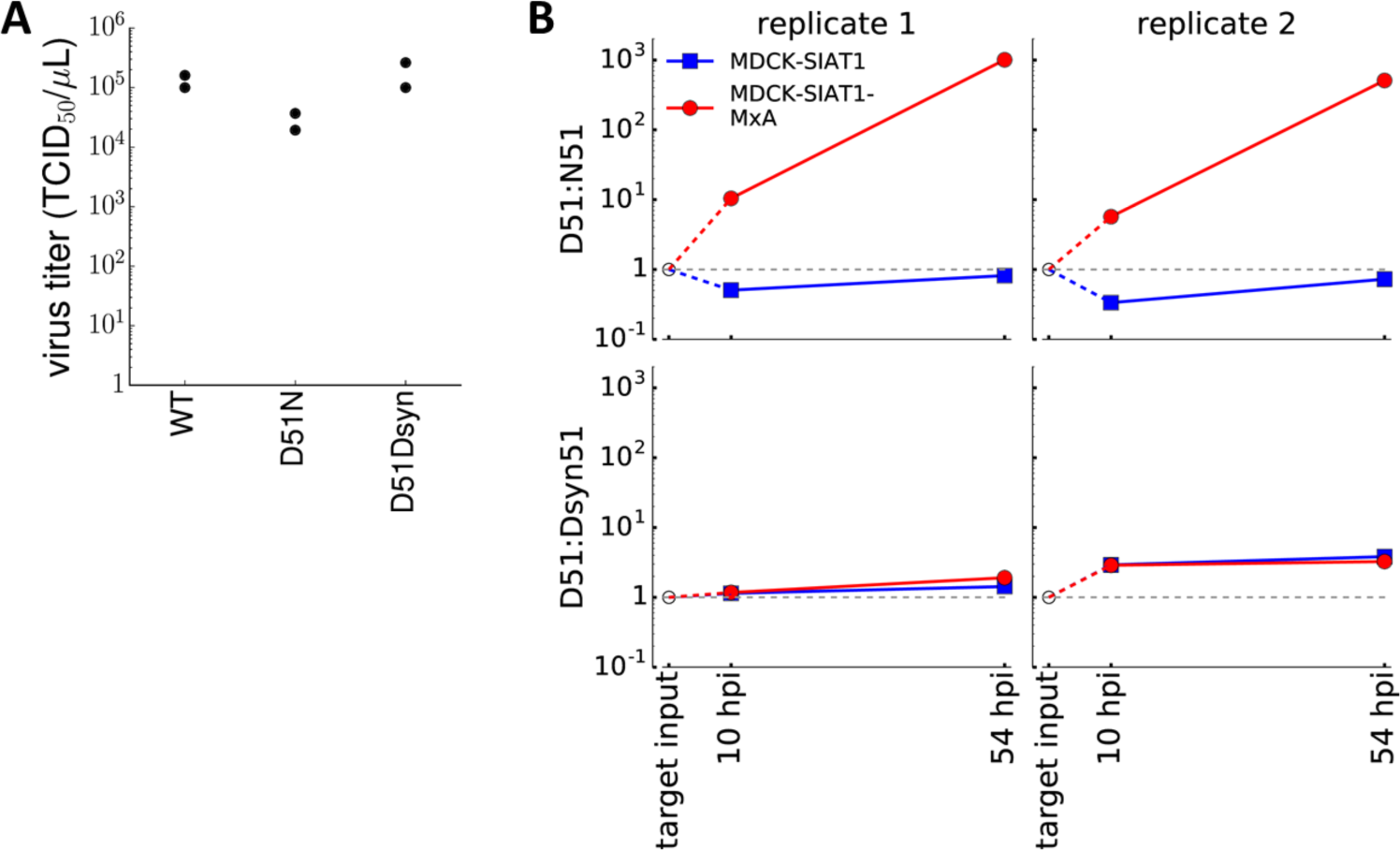
A mutation a site 51 in NP increases MxA sensitivity as measured by viral competition. (**A**) We used reverse genetics to generate viruses carrying the wildtype NP, the D51N mutation, or a synonymous mutation at site 51. Each virus was generated in duplicate. The plot shows that all viruses reached titers that were within 5-fold of each other after 72 hours. (**B**) We mixed the wildtype virus with one of the mutant viruses, infected MxA-expressing and non-expressing cells, and measured mutant frequencies after 10 and 54 hours. The plots show the frequency of the wildtype D51 variant relative to the N51 amino-acid mutant or the Dsyn51 synonymous mutant at each timepoint. In both replicates of the competition, the wildtype D51 variant was strongly favored over the N51 mutant in MxA-expressing cells. We targeted a 1:1 ratio of infectious particles in our initial inoculum, but this ratio was impossible to verify by direct sequencing since sequencing cannot distinguish infectious from non-infectious virions.

## Discussion

We have used deep mutational scanning to experimentally estimate how MxA sensitivity is affected by every amino-acid mutation to a NP from a human-adapted influenza strain. Our approach screens all possible mutations, and can identify changes that increase or decrease MxA resistance. In contrast, previous approaches have focused on mutations that fixed during viral adaptation to MxA in nature or in the lab [17, 22].

These approaches have different strengths. Examining mutations that fix during viral adaptation elucidates the evolutionary pathways to MxA resistance. Our approach systematically maps how all sites in NP contribute to MxA resistance, without regard to whether mutating these sites in the starting virus can increase MxA resistance. The two approaches will yield similar results if all sites start with amino acids that confer sensitivity to MxA. But the approaches will yield different results if most sites in the starting virus already have amino acids that confer resistance to MxA.

The most striking finding of our work is that the latter situation predominates: most sites with the largest effect on MxA resistance already possess amino acids that confer resistance. Furthermore, at most of these sites, the resistant amino acid is conserved across human and avian influenza strains. As a consequence, most sites that we identified could not have been found by looking for mutations that adapt influenza to MxA in nature or the lab for the simple reason that these sites are already fixed to a resistant amino acid.

Why are so many sites in NP already fixed at MxA-resistant identities? One possibility is that homologs of MxA in other hosts have selected for some level of generalized MxA resistance in all NPs. Determining whether this is the case will require characterizing whether MxA homologs of the relevant species do in fact exert selection on influenza: there is evidence that swine MxA restricts influenza [46, 47], but the anti-influenza activity of avian MxA remains incompletely characterized across most bird species that are hosts for influenza [48–50]. Even if other MxA homologs exert selection, we would not expect non-human viruses to be optimally resistant to human MxA since MxA homologs have different specificities [51]. But similarities among the recognition mechanisms of MxA homologs could have driven fixation of resistant amino acids at many sites. Alternatively, perhaps many sites in NP have MxA-resistant amino acids due to some unknown selective pressure unrelated to MxA. In any case, our results demonstrate that it is important to establish the baseline when thinking about MxA resistance. While avian influenza strains are more sensitive to human MxA than human strains [16, 17], our results suggest that these avian strains still have amino acids that confer MxA resistance at most sites.

Another question is whether MxA resistance comes at an inherent functional cost. Several studies have introduced resistance mutations from human NPs into avian NPs and found that the resulting viruses are attenuated [17, 45]. One interpretation is that MxA resistance is inherently costly. But another interpretation is simply that amino-acid mutations are often detrimental, and that this is no more likely to be true of MxA-resistance mutations than MxA-sensitizing ones. In support of this idea, the MxA-sensitizing mutations that we identified at site 51 were all slightly deleterious to viral polymerase activity. In addition, the functional effects of mutations affecting MxA resistance may sometimes be idiosyncratic to the particular viral strain. For instance, the MxA-sensitizing D51N mutation was found to be beneficial for viral growth in an engineered avian influenza NP [45], but was deleterious to both polymerase activity and viral growth in the Aichi/1968 NP that we used in our study.

A related question is whether mutations that affect MxA resistance in one NP similarly affect resistance in NPs from more diverged viral strains. Our results suggest that the answer is yes. For instance, reverting each of the three mutations responsible for the MxA resistance of the 1918 virus [17] led to the expected decrease in MxA resistance in the Aichi/1968 NP used in our study. Similarly, our finding that D51N greatly increases MxA sensitivity agrees with another study [45] that reported this mutation also increased the MxA sensitivity of an engineered avian NP. Therefore, it appears that mutational effects on MxA sensitivity are at least somewhat conserved across NPs.

We also find great consistency in the regions of NP where mutations affect MxA sensitivity. Despite the fact that most sites that we identified as contributing to MxA resistance are new, many of these sites map to the same regions of NP as previously characterized resistance mutations. About half the sites that we identified clustered at the base of the NP body domain, which is also the location of resistance mutations in the 1918 and 2009 pandemic viruses, as well as H7N9 influenza [17, 22]. This solvent-exposed region, which is distinct from the NP surfaces important for RNA binding or interactions with the polymerase proteins [52, 53], could plausibly be a binding interface between MxA and NP. However, we also found that MxA sensitivity was affected by some sites in surface-exposed loops distal from the base of the NP body domain. So although our results confirm that certain regions of NP are disproportionately important determinants of MxA sensitivity, the details of the inhibitory interaction between NP and MxA remain unclear [10, 54].

Overall, we have used a powerful new deep mutational scanning approach to identify sites that affect the interactions between a restriction factor and its viral target. This approach complements the traditional strategy of characterizing mutations at specific sites that differ across viral strains. An advantage of our approach is that it enables unbiased identification of all sites where mutations affect a phenotype, regardless of whether these sites have substituted during evolution. We envision that this approach can be extended to systematically examine how viral mutations affect additional homologs of NP and MxA, as well as to understand the interactions between viruses and other less well-characterized restriction factors [55, 56].

## Materials and Methods

### Availability of data and computer code

FASTQ files for the deep mutational scanning experiment and viral competition experiment are on the NCBI Sequence Read Archive with accession number SRP082554. The computer code and input data files necessary to reproduce all the analysis in this work are available inS1 File and also at https://github.com/orrzor/2016_NP_MxA_paper (last accessed August 15, 2016). The differential selection values estimated at each site in NP, and for each mutation at each site in NP, are inS2 File andS3 File respectively.

### Plasmids

#### Plasmids for polymerase activity assays

The polymerase activity assays used a plasmid that transcribed a reporter viral RNA expressing GFP with flanking regions from the PB1 gene from the A/WSN/1933 (H1N1) strain. A version of this reporter driven off a human RNA polymerase I promoter has been previously described as pHH-PB1flank-eGFP [57]. The experiments here utilized variants of the canine MDCK cell line, so we needed to clone this reporter into a plasmid with the canine RNA polymerase I promoter. Based on the description of this promoter provided in [58], we created a recipient plasmid for BsmBI-based cloning of viral RNAs under control of a canine RNA polymerase I promoter, and named this plasmid pICR2 (plasmid influenza canine reporter 2); a map of this plasmid is inS4 File. We then cloned the GFP-containing reporter from pHH-PB1flank-eGFP into this plasmid to create pICR2-PB1flank-eGFP; a map of this plasmid is inS5 File.

The NP protein for these polymerase activity assays was expressed from the pHW-Aichi68-NP plasmid described in [59] that encodes the NP from human influenza strain A/Aichi/2/1968 (H3N2), or from variants of this plasmid constructed by site-directed mutagenesis that expressed the point mutants of the NP described in the current paper. The polymerase proteins were derived from the human influenza strain A/Nanchang/933/1995 (H3N2), and were expressed from plasmids HDM-Nan95-PB2, HDM-Nan95-PB1, or HDM-Nan95-PA. The HDM plasmid expresses genes off a CMV promoter, and the PB2 / PB1 / PA gene sequences themselves are those given in [59].

#### Plasmids for influenza reverse genetics

The viruses used in these experiments had the same composition as those used in [59]: the NP was from A/Aichi/2/1968 (plasmid pHWAichi68-NP), the three polymerase genes were from A/Nanchang/933/1995 (plasmids pHWNan95-PB2, pHWNan95-PB1, and pHWNan95-PA), and the remaining genes were from A/WSN/1933 (plasmids pHW184-HA, pHW186-NA, pHW187-M, and pHW188-NS). The reverse genetics plasmids for the first four of these genes are described in [59], while the last four are described in [60]. The mutants of the NP used in this study were created by introducing point mutations into pHW-Aichi68-NP.

### Cell lines

We used lentiviral transduction to engineer MDCK-SIAT1 cells to constitutively express human MxA or MxA-T103A under control of a CMV promoter. We placed a FLAG tag followed by a GSG linker (DYKDDDDKGSG) after the methionine start codon of human MxA. Downstream of MxA, we included an internal ribosome entry site (IRES) followed by the red fluorescent protein mCherry to act as a reporter for lentiviral transduction. At 48 hours after lentiviral transduction of the MDCK-SIAT1 cells, we single-cell cloned variants by serial dilution in 96-well plates. Wells with clonal transduced cells were identified by finding wells with single clusters of cells expressing mCherry.

To verify that the recovered cell lines expressed MxA, we seeded the cells at 2.5 × 10^5^ cells/well in D10 media (DMEM supplemented with 10% heat-inactivated FBS, 2 mM L-glutamine, 100 U of penicillin/ml, and 100 *µ*g of streptomycin/ml) in 12-well dishes, and 20 h later, we collected the cells and performed Western blots to detect the FLAG tag on the MxA or β-actin as a loading control. To detect FLAG, we stained with a 1:5000 dilution of mouse anti-FLAG (Sigma, F1804) followed by a 1:2500 dilution of Alexa Flour 680-conjugated goat anti-mouse (Invitrogen, A-21058). To detect *β*-actin, we stained with a 1:5000 dilution of rabbit anti-*β*-actin (Abcam, ab8227) followed by a 1:2000 dilution of Alexa Flour 680-conjugated goat anti-rabbit (Invitrogen, A-21109). Blots were imaged using a Li-Cor Odyssey Infrared Imaging System.

### Deep mutational scanning

#### Experimental selections in MxA expressing and control cell lines

Our deep mutational scanning used the duplicate Aichi/1968 mutant viral libraries that had been passaged in MDCK-SIAT1 cells described in [32]. These viral libraries carried mutant Aichi/1968 NP, PB1/PB2/PA from A/Nanchang/933/1995 (H3N2) and HA/NA/M/NS from A/WSN/1933 (H1N1). Viruses were grown in the WSN growth media described in [30] (Opti-MEM supplemented with 0.5% heat-inactivated FBS, 0.3% BSA, 100 U of penicillin/ml, 100 *µ*g of streptomycin/ml, and 100 *µ*g of calcium chloride/ml). Trypsin was not added to the media as viruses with the WSN/1933 HA and NA are trypsin independent [61].

The seven samples for each deep mutational scanning replicate are shown in S1 Fig. The*DNA*, *mutant DNA*, *virus*, and*mutant virus* samples have been described previously [32]. The remaining three samples (*mutant virus MDCK-SIAT1-MxA*, *mutant virus MDCK-SIAT1*, and*mutant virus MDCK-SIAT1-MxA-T103A*) were prepared as follows: the appropriate MDCK-SIAT1 cell line variant was plated in D10 media in a 10-cm dish at 3.2 × 10^6^ cells/dish. After 14 hours, the media was changed to WSN growth media containing mutant virus library diluted so that the MOI of infection was 0.1 TCID_50_ per cell. Each mutant virus was used to infect eight 10-cm dishes for a total of 4.8 × 10^6^ TCID_50_ passaged per selective condition (the cells have increased to 6 × 10^6^ / dish by the time of infection). After 2 hours, the media was changed to fresh WSN growth media. At 48 hours post-infection, the viral supernatant was collected, clarified by centrifugation at 2, 000 × *g*, and stored at 4^*^o^*^C. For each sample, virus was pelleted by centrifuging 25 mL of clarified viral supernatant at 64, 000 × *g* for 1.5 h at 4^*^o^*^C. Viral RNA was then extracted and prepared for sequencing on an Illumina HiSeq 2500 using 150-bp paired end reads in rapid run mode as described in [32].

#### Counting mutations from deep sequencing data

To obtain higher sequencing accuracy, we mapped reads so that we only counted codon identities at sites covered by overlapping paired-end reads, requiring that both reads in a pair concurred at a site. This mapping was done using mapmuts (https://github.com/jbloom/mapmuts; version 1.1) as described in [32]. The result of this mapping is a count of the number of each codon (and amino acid) identity at each site in NP for each sample.

#### Quantifying differential selection from mutation counts

In order to quantify differential selection from the counts of mutations in each sample, we developed a metric for differential selection that is loosely based on the notion of selection coefficients described in [62]. Briefly, if some non-wildtype amino-acid *a* at site *r* is observed 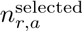 times in the selected sample and 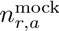 in the mock-selected sample, then we compute the relative enrichment *E*_*r*,*a*_ of *a* at site *r* as 

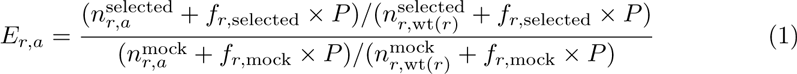
where wt (*r*) denotes the wildtype amino acid at site *r*, *P* is a pseudocount (set to 10 in our analyses), *f*_*r,*_ selected and *f*_*r*,_ mock give the relative depths of the selected and mock samples:

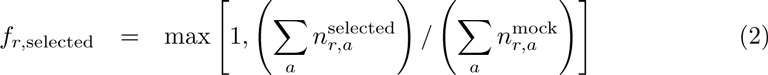

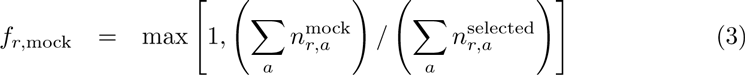

The reason for scaling the pseudocount by the library depth is that in the absence of such scaling, if the selected and mock samples are sequenced at different depths, the estimates of *E*_*r*,*a*_ will tend to be systematically different from one even if there the relative counts are the same in both conditions. The differential selection values are the logarithm of the enrichment values:

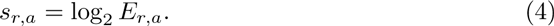

It is these differential selection values that are reported in the manuscript. For the MxA selections, the values are computed by comparing the counts in the MxA-expressing cells to those in the non-expressing cells. For the control selections, the values are computed by comparing the counts in the cells expressing the inactive T103A MxA to those in the non-expressing cells.

Computer programs to calculate the differential selection values and display them in logo plots were implemented into the dms tools software [63] available at https://github.com/jbloomlab/dms_tools. We used version 1.1.17 of this software for the analyses in this paper.

### Viral polymerase activity

We measured polymerase activity for different NP mutants in the MDCK-SIAT1 cells expressing or not expressing MxA. For these assays, we co-transfected 500 ng of the pICR2-PB1flank-eGFP reporter plasmid along with 15 ng of indicated mutant pHW2000-Aichi68 NP and 125 ng each of HDM-Nan95-PA, HDM-Nan9-PB1, and HDM-Nan95-PB2 into wells of 24-well dishes of MDCK-SIAT1 or MDCK-SIAT1-MxA cells. We chose this amount of NP plasmid because it was near the midpoint of the polymerase activity dose-response curve when holding the amounts of all other plasmids fixed (S6 Fig).

Transfections were performed with Lipofectamine 3000 (ThermoFisher Scientific) and the transfection mixes of DNA and lipofectamine were prepared according to the manufacturer’s protocol. The transfection itself was done using the modified protocol below, which was designed to increase transfection efficiency in MDCK-SIAT1 cells [64]. The transfection mix was incubated at room temperature for 1 hour and added to a well of a 24-well plate, and then 500 *µ*L cells at 2.5 × 10^5^ cells/mL was added to this well. After 4 hours, we changed the media to fresh D10. At 20 hours post-transfection, we quantified the geometric mean of the GFP fluorescence by flow cytometry. We performed three biological replicates for each NP mutant, and each replicate used NP plasmid from an independent mini-prep.

### Viral competition

#### Generating mutant viruses

The viruses used in the competitions had the same composition as those in the deep mutational scanning: NP from A/Aichi/2/1968, polymerase genes from A/Nanchang/933/1995, and the remaining genes from A/WSN/1933. These viruses were generated by reverse genetics [60] using pHWAichi68-NP, pHWNan95-PB2, pHWNan95-PB1, pHWNan95-PA, pHW184-HA, pHW186-NA, pHW187-M, and pHW188-NS. The viruses were titered in MDCK-SIAT1 cells using the TCID_50_ protocol described in [28].

#### Competitions

For each viral competition, MxA-expressing and non-expressing MDCK-SIAT1 cells were infected with a mixture of wildtype and mutant virus in duplicate. Virus carrying wildtype Aichi/1968 NP and virus carrying a mutant Aichi/1968 NP were mixed in a 1:1 ratio based on TCID_50_. MDCK-SIAT1 cells and MDCK-SIAT1-MxA cells were plated in D10 media in 6-well dishes at 2.5 × 10^5^ cells/mL. We infected cells in WSN growth media at MOIs of 0.1 and 0.01 for the 10 hour and 54 hour timepoints, respectively. At 2 hours post-infection, media was changed to new WSN growth media. At 10 hours and 54 hours post-infection, cellular RNA was extracted using the Qiagen RNeasy Plus Mini kit. Cells were lysed in buffer RLT Plus supplemented with β-mercaptoethanol and RNA was extracted following the manufacturer’s protocol.

#### Sequencing to determine mutant frequency

We reverse transcribed the NP gene from the extracted cellular RNA using SuperScript III Reverse Transcriptase (ThermoFisher Scientific) following the manufacturer’s protocol for first-strand cDNA synthesis. The reverse transcriptase reaction contained 500 ng cellular RNA template and used the primers 5’-BsmBI-Aichi68-NP (5’-CATGAT CGTCTCAGGGAGCAAAAGCAGGGTAGATAATCACTCACAG-3’) and 3’-BsmBI-Aichi68-NP (5’-CATGATCGTCTCGTATTAGTAGAAACAAGGGTATTTTTCTTTA-3’) to collect both positive-sense and negative-sense viral RNA templates.

We then carried out targeted deep sequencing of the mutated region in NP [65]. We first amplified a region of the NP gene centered around codon site 51 using 2 *µ*L cDNA template and the primers 5’-CTTTCCCTACACGACGCTCTTCCGATCTGATTCTACATCCAAATGTGCACTGAACTTAAAC-3’ and 5’-GGAGTTCAGACGTGTGCTCTTCCGATCTTGTTAAGCTGTTCTGGATCAGTCGC-3’, which added a part of the Illumina sequencing adaptors. We used the following PCR program.

1. 95 °C for 2 min.
2. 95 °C for 20 s.
3. 70 °C for 1 s.
4. 55 °C for 10 s.
5. 70 °C for 20 s.
6. Repeat steps 2 through 5 for 24 additional cycles.
7. Hold 4 °C.

We purified the PCR product using 1.5X AMPure XP beads (Beckman Coulter) and used 5 ng of this product as template for a second round of PCR using the following pair of primers that added the remaining part of the Illumina sequencing adaptors as well as a six-mer barcode (xxxxxx) used to differentiate the experimental samples: 5’-A ATGATACGGCGACCACCGAGATCTACACTCTTTCCCTACACGACGCTCTTC CGATCT-3’ and 5’-CAAGCAGAAGACGGCATACGAGATxxxxxxGTGACTGGAGT TCAGACGTGTGCTCTTCCGATCT-3’. We used the PCR program below.

1. 95 °C for 2 min.
2. 95 °C for 20 s.
3. 70 °C for 1 s.
4. 58 °C for 10 s.
5. 70 °C for 20 s.
6. Repeat steps 2 through 5 for 4 additional cycles.
7. Hold 4 °C.

All experimental samples were then pooled, gel purified, and sequenced with 75-bp paired-end reads using an Illumina MiSeq with the MiSeq Reagent Kit v3 150-cycle.

We processed the sequencing reads to determine the frequency of the mutant NP virus in each competition. Both reads in each read pair were first filtered. Any position with Q-score below 15 was assigned as the ambiguous nucleotide N, and if a read had more than 5 ambiguous nucleotides, it was discarded. Next, both reads in each read pair were aligned to the Aichi/1968 NP. Sites overlapping between reads were counted as ambiguous unless both reads agreed at the site. Read pairs were discarded if there were more than 2 codon mutations. Finally the read pair was translated and the numbers of wildtype and mutant amino acids at codon site 51 were counted.

## Acknowledgments

We thank the Fred Hutch Genomics Core for performing the deep sequencing, and the Fred Hutch Flow Cytometry Core facilities for assistance with flow cytometry. We thank Michael Emerman, Harmit Malik, Adam Dingens, Hugh Haddox, Alistair Russell and Shirleen Soh for comments on the manuscript.

## Supporting Information

**S1 Fig.**
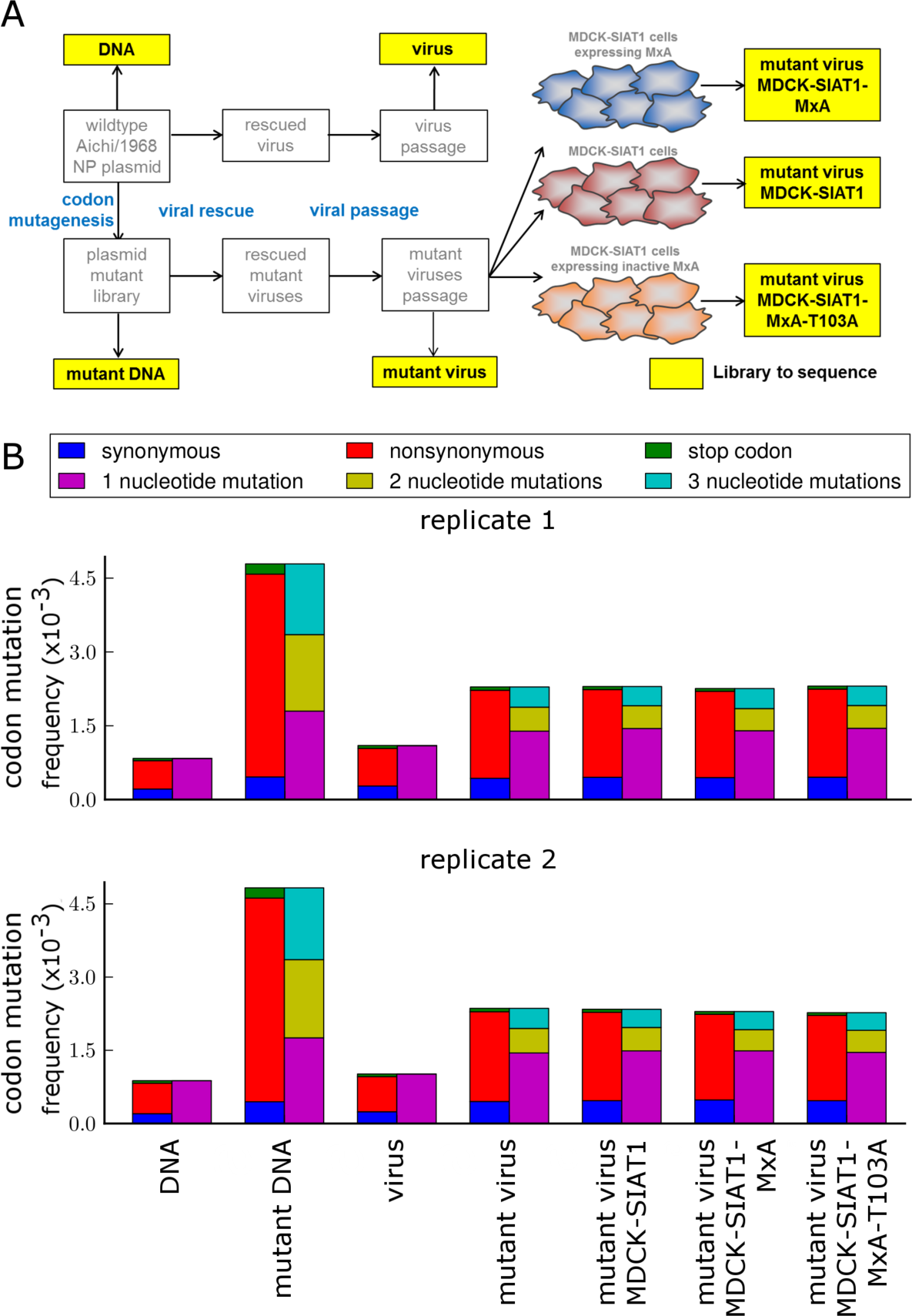
Mutation and error rates of deep sequenced libraries. (**A**) Schematic of samples sequenced. Beginning with a plasmid carrying unmutated NP, we created a library of plasmids encoding codon mutants of NP as described in [32]. We then used reverse genetics to generate wildtype and mutant viruses from unmutated and mutagenized plasmids and passaged these viruses at low MOI in MDCK-SIAT1 cells as described in [32]. We then infected the mutant virus libraries into the three cell lines shown in Fig. 1A. (**B**) The average per-codon mutation frequency for each sample in each library. Codon mutations are classified by the type of mutation or the number of nucleotide changes relative to the wildtype codon. Mutation rates for the mutant virus passaged through MxA-expressing cells and for mutant virus passaged through MxA non-expressing or inactive cells were similar. However this does not preclude interesting MxA selection occurring on a site-by-site basis. Note that the background error rates estimated from sequencing the unmutated DNA and virus are less than the mutation rates in the libraries; in addition, these errors affect only the single-nucleotide codon mutations.

**S2 Fig.**
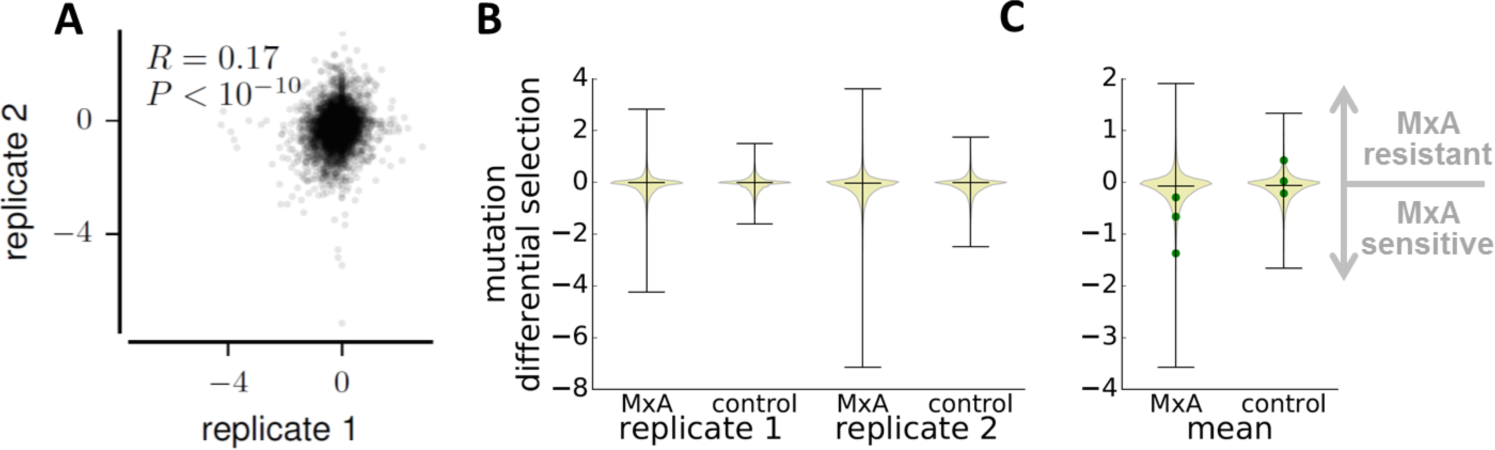
Differential selection on individual mutations across all NP sites. This figure differs from Fig.3 by showing data for individual mutations rather than sites (see Fig. 1B for the distinction between mutation and site differential selection). (**A**) Differential selections for individual mutations are weakly correlated between experimental replicates. The weakness of this correlation relative to that for the site differential selections shown in Fig 3C is likely because at any NP site that is a determinant of MxA resistance, several mutations usually affect MxA resistance. Therefore even if replicates do not evenly sample the same individual mutations at that site [30, 32], both replicates will still detect similar total differential selection when averaged across all mutations at a site. (**B**) Distributions of differential selection for all mutations. Distributions are displayed for the actual (MDCK-SIAT1-MxA vs MDCK-SIAT1) and control (MDCK-SIAT1-MxA-T103A vs MDCK-SIAT) selections for both replicates of the mutational scanning. The selection distributions have longer tails than the control distributions, which indicates that some mutations affect MxA sensitivity more than the background experimental noise. (**C**) The distributions of mutation differential selections for the mean of the replicates. The differential selection values for mutations V100R, P283L, and Y313F previously demonstrated [17] to increase MxA sensitivity in 1918 virus are shown in green. Each mutation increased MxA sensitivity in our data as expected.

**S3 Fig.**
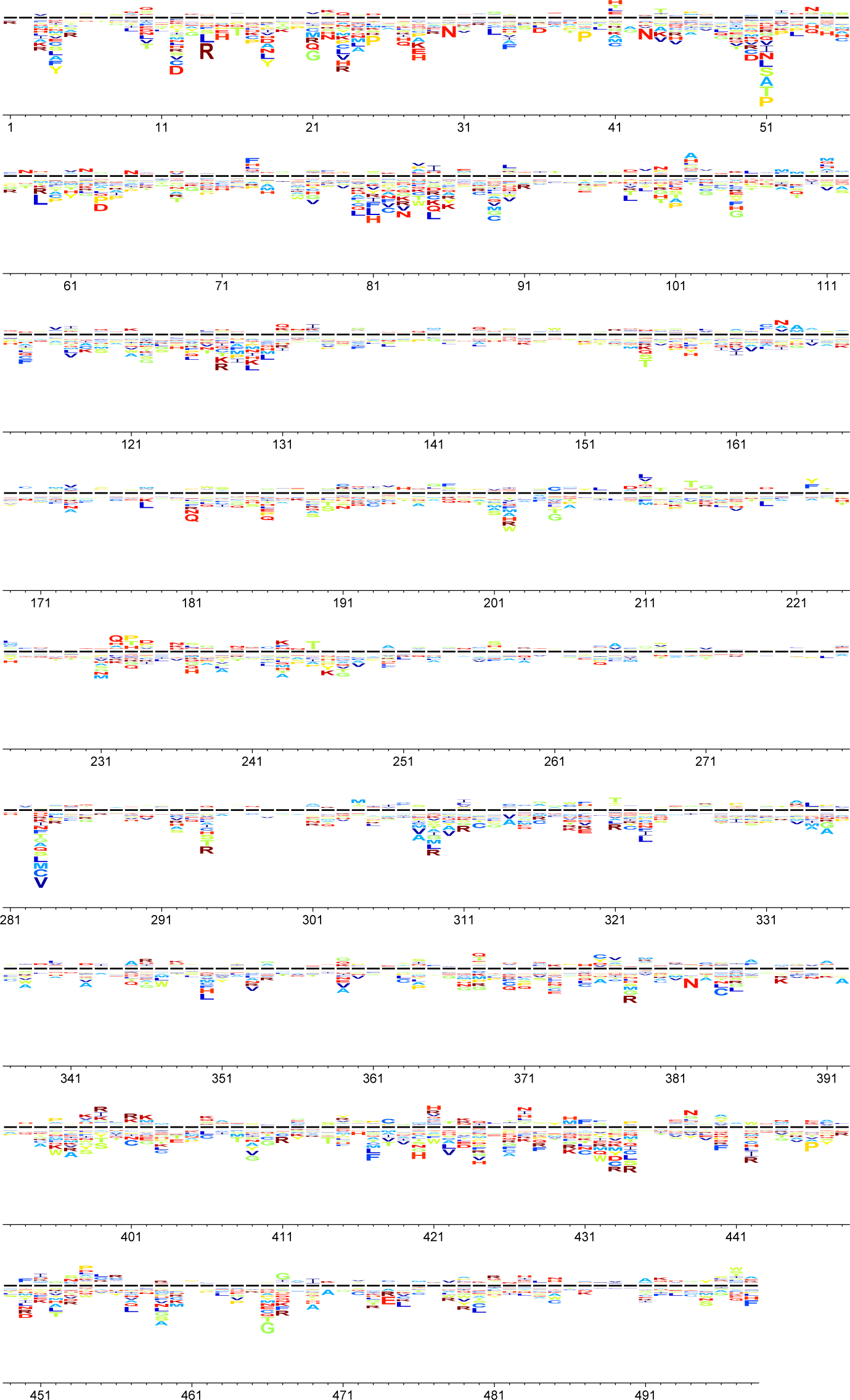
Differential selection from MxA at each site in NP. The height of each letter above or below the black center line indicates selection for or against that mutation in MxA-expressing versus non-expressing cells (see Fig. 1B). These data are the average of the two replicates, and the values are given inS3 File. Letters are colored by amino-acid hydrophobicity.

**S4 Fig.**
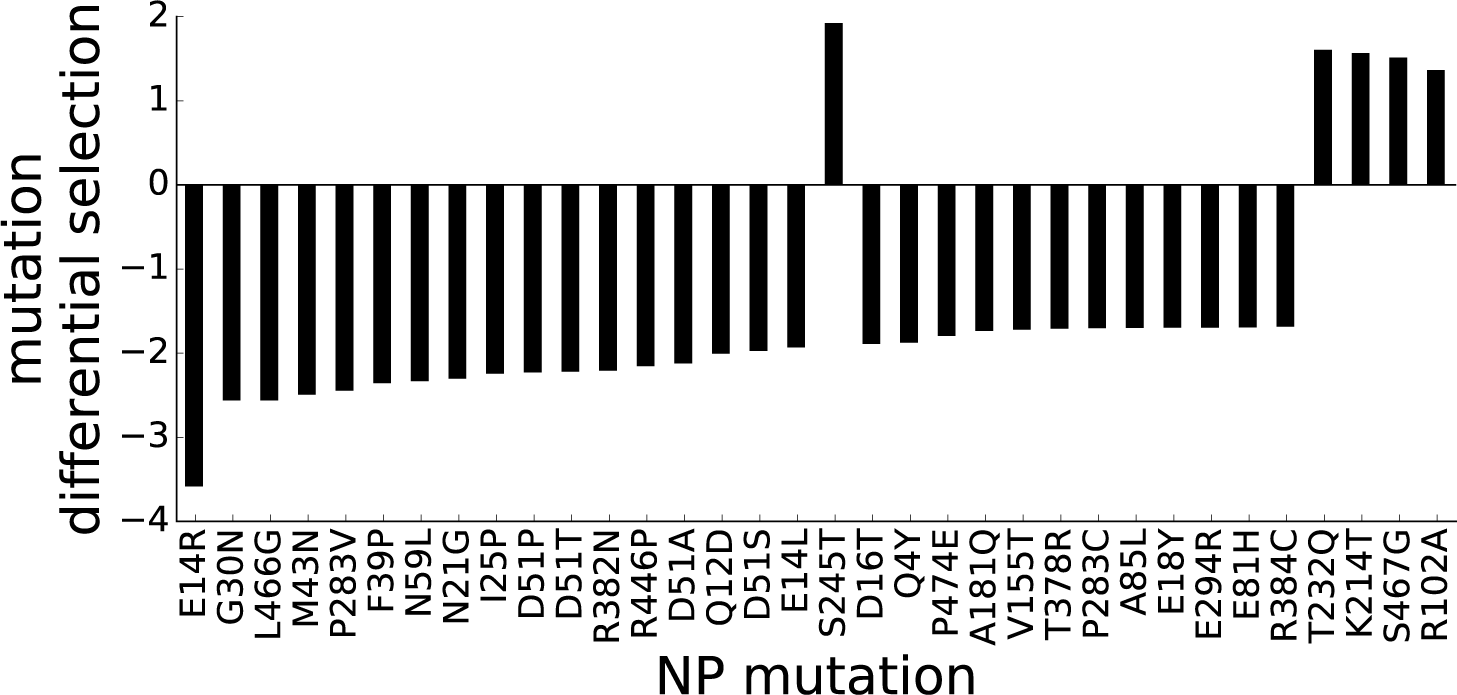
Most NP mutations with the greatest differential selection increase MxA sensitivity. Differential selections for mutations in MxA-expressing vs non-expressing cells after averaging the data for the two replicates. Shown are all individual mutations that have differential selections more extreme than that seen in the control selections: 5 mutations have differential selections greater than the maximum seen in the control selection, and 29 mutations have differential selections less than the minimum seen in the control selection. Most mutations have negative differential selection, and 14 of 34 mutations occur at the sites shown in Fig 4B as having high differential selection.

**S5 Fig.**
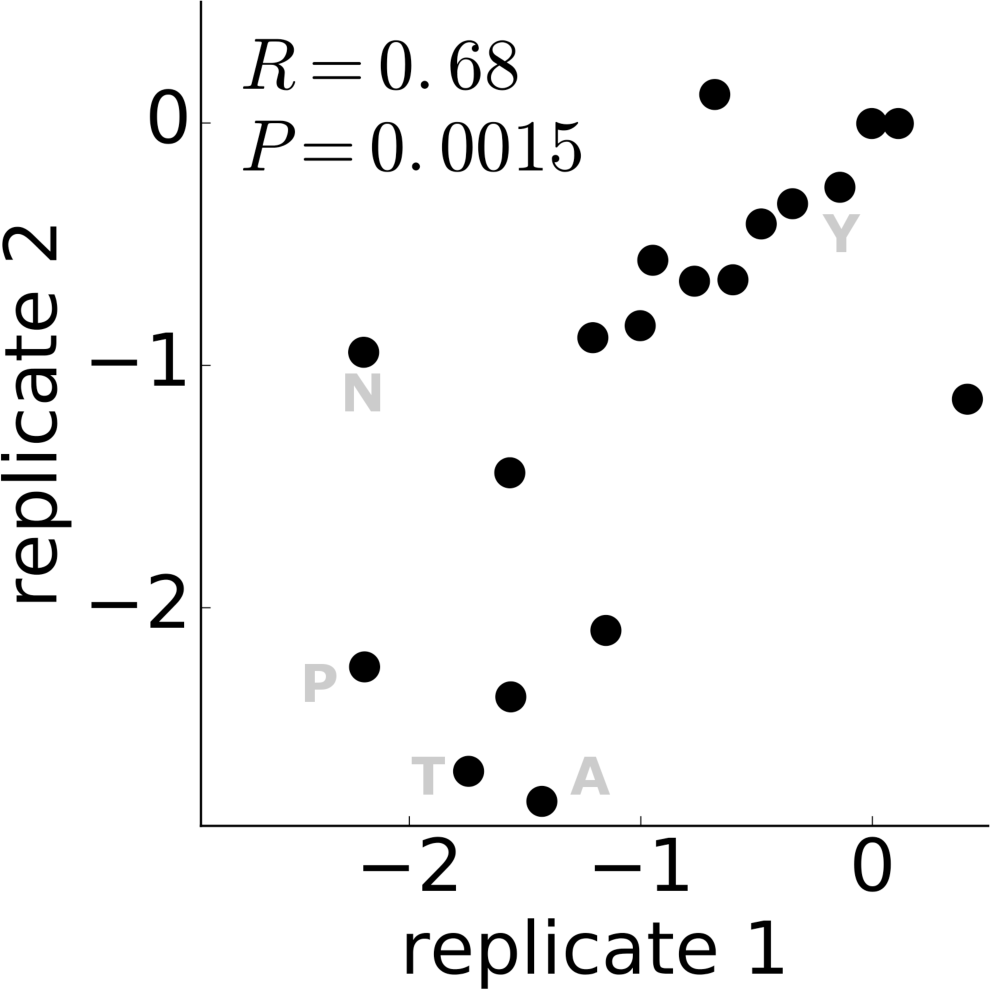
Differential selection of mutations at site 51 in NP. The differential selection for MxA resistance for the mutations at site 51 for both replicates of the deep mutational scanning. A negative value indicates the mutation increases MxA sensitivity.

**S6 Fig.**
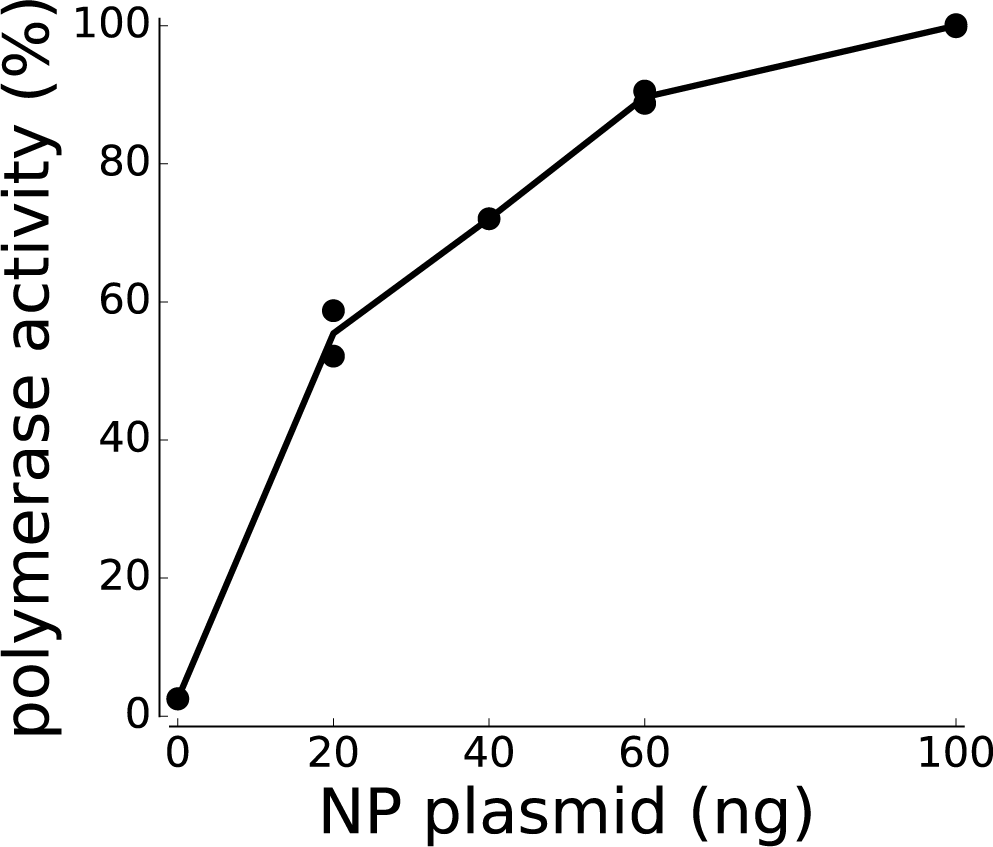
Determining the amount of NP to use in the polymerase activity assay. The polymerase activity as a function of the amount of transfected NP plasmid. The Aichi/1968 NP plasmid was varied from 0 to 100 ng, while the PB1, PB2, and PA plasmids were fixed at 125 ng each and the GFP reporter plasmid was fixed at 500 ng. The experiment was done in duplicate and the polymerase activity of the mix with the maximum amount of NP is set to 100%. For all our polymerase activity assays, we chose an amount of NP (15 ng) near the midpoint of the dose-response curve. At this level of NP, the reporter signal is sensitive to changes in the amount of NP due to MxA activity.

**S1 Table.**
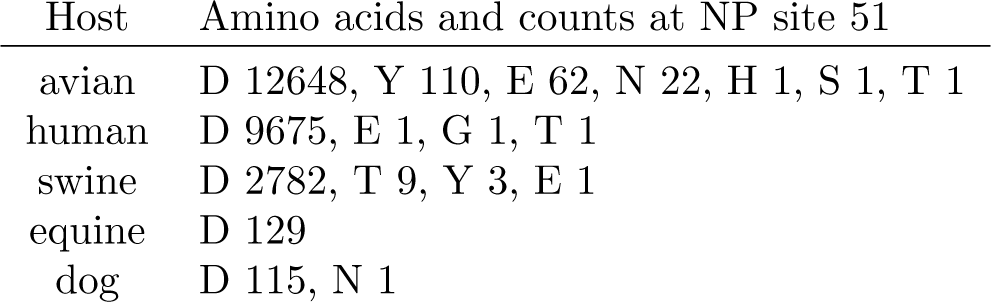
Amino-acid counts at NP site 51 across influenza hosts.

| Host | Amino acids and counts at NP site 51 |
| --- | --- |
| avian | D 12648, Y 110, E 62, N 22, H 1, S 1, T 1 |
| human | D 9675, E 1, G 1, T 1 |
| swine | D 2782, T 9, Y 3, E 1 |
| equine | D 129 |
| dog | D 115, N 1 |

For influenza strains from each host, we downloaded all available full-length NP protein sequences from the Influenza Research Database (https://www.fludb.org) (last accessed August 15, 2016). Across all strains, D occurs at site 51 with 99% frequency.

**S1 File. Computer code and data.** This ZIP file contains the computer code and input data files to perform the analysis described in this paper.

**S2 File. Differential** selection values for all sites. This text file contains the estimated differential selection at each site in NP after averaging the measurements across replicates.

**S3 File.** Differential selection values for all mutations. This text file contains the estimated differential selection on each mutation at each site in NP after averaging the measurements across replicates.

**S4 File.** Map of pICR2, the recipient plasmid with the canine RNA polymerase I promoter. The map is provided as a Genbank file.

**S5 File.** Map of pICR2-PB1flank-eGFP, the GFP reporter plasmid with canine RNA polymerase I promoter. The map is provided as a Genbank file.

